# The regulatory landscapes of human ovarian ageing

**DOI:** 10.1101/2022.05.18.492547

**Authors:** Chen Jin, Xizhe Wang, Adam D. Hudgins, Amir Gamliel, Mingzhuo Pei, Seungsoo Kim, Daniela Contreras, Jan Hoeijmakers, Judith Campisi, Rogerio Lobo, Zev Williams, Michael G. Rosenfeld, Yousin Suh

## Abstract

The ovary is the first organ to age in the human body, affecting both fertility and overall health in women^1-8^. However, the biological mechanisms underlying human ovarian ageing remain poorly understood. Here we performed single-nuclei multi-omics analysis of young and reproductively aged human ovaries to understand the molecular and cellular basis of ovarian ageing in humans. Our analysis reveals coordinated changes in transcriptomic output and chromatin accessibility across cell types during ageing, including elevated mTOR and MAPK signaling, decreased activity of the oxidative phosphorylation and DNA damage repair pathways, and an increased signature of cellular senescence. By constructing cell type-specific regulatory networks, we uncover enhanced activity of the transcription factor CEBPD across cell types in the aged ovary, with a corresponding significant loss of activity of most cell identity-associated transcription factors. Moreover, by performing integrative analyses of our single-nuclei multi-omics data with common genetic variants associated with age at natural menopause (ANM) from genome-wide association studies, we demonstrate a global impact of functional variants on changes in gene regulatory networks across ovarian cell types. Finally, we nominate about a dozen of functional non-coding variants, their target genes and cell types and regulatory mechanisms that underlie genetic association with ANM. This work provides a comprehensive multimodal landscape of human ovarian ageing and mechanistic insights into inherited variation of ANM.

## Main

The ovary is the primary female reproductive organ, and the first tissue to undergo profound age-associated loss of function in humans, characterized by a progressive decline in follicle number and oocyte quality^1^. The rate of follicular depletion increases throughout reproductive life, but begins a more accelerated decline around age 37^9^. This results in a higher risk of both infertility, and aneuploidy and congenital disabilities in offspring^2^. There is also overwhelming evidence that female reproductive ageing influences lifespan and diverse health outcomes^3-8,10,11^. Consequently, an in-depth understanding of ovarian ageing can benefit not only reproduction but also longevity and health in women. However, thus far very little is known about basic biological mechanisms that underlie human ovarian ageing.

Menopause is the time marking the cessation of menstrual cycling and production of fertile oocytes, and age at natural menopause (ANM) has profound implications for health and disease risk in women^3-8^. Family and twin studies have demonstrated a strong relationship between genetics and ANM^12-15^, suggesting up to a ∼6-fold increase in risk of early menopause for a woman with a family history of early menopause^12,15^. Identification of the genes contributing to ANM will provide mechanistic insights into the biological processes underlying ovarian ageing. Genome-wide association studies (GWAS) have identified hundreds of genetic loci associated with ANM^16^. However, the great majority (∼94%) of the risk variants reside in non-coding regions of the genome, making it difficult to assign their functional role in ovarian ageing

Many recent studies show that functional non-coding GWAS variants are significantly enriched in cell type-specific transcriptional regulatory elements such as enhancers^17-25^. Enhancers have emerged as major points of integration of intra- and extracellular signals associated with development, homeostasis, and disease, resulting in context-specific transcriptional outputs^26^. Cell-specific enhancer activation is driven by combinatorial actions of lineage-determining and signal-dependent transcription factors (TFs)^27^. Genetic variation affecting enhancer selection and function is considered a major determinant of differences in cell-specific gene expression and disease risk between individuals^27^. Therefore, the identification of functional ANM-associated non-coding regulatory variants, as well as the target genes and cell types through which they confer their effects on ANM, is a powerful way to understand the biological processes underlying ovarian ageing. However, we currently lack an atlas of the transcriptional regulatory elements that are active in every cell type in the ovary during ageing.

In this study, we systematically characterize human ovarian ageing by performing single-nuclei multi-omics analysis and by superimposing these data with ANM-associated GWAS risk variants. Through these efforts, we identify the functional transcriptional regulatory elements, and functional non-coding variants and their target genes associated with ANM, across all cell types in the ovary.

### Single nucleus multi-omics profiling ageing

We performed single-nuclei RNA-seq (snRNA-seq) and single-nuclei assay for transposase-accessible chromatin using sequencing (snATAC-seq) on the same flash-frozen human ovarian tissues, which were from young (n=4; ages 23-29 years) and reproductively old (n=4; ages 49-54 years) autopsy samples of sudden death with normal ovarian histology (Fig. 1a and Supplementary Table 1). After stringent quality control, we retained 42,568 nuclei for snRNA-seq and 41,550 nuclei for snATAC-seq (Methods). Seurat-based unsupervised clustering^28^ and Harmony-based^29^ batch correction on snRNA-seq revealed eight distinct clusters (Methods, Fig. 1b, and Supplementary Fig. 1a). All major somatic cell types in the ovary, including stromal cells (SC), endothelial cells (blood vessel endothelial cells (BEC) and lymphatic endothelial cells (LEC)), granulosa cells (GC), smooth muscle cells (SMC), immune cells (IC), epithelial cells (EpiC), and theca cells (TC) were identified based on well-defined cell type-specific markers (Figs. 1b, c). For snATAC-seq, Signac-based unsupervised clustering^30^ and Harmony-based batch correction revealed seven distinct clusters (Methods, Fig. 1d, and Supplementary Fig. 1b). To annotate the clusters, we used canonical correlation analysis (CCA) and mutual nearest neighbors (MNNs)^31^ to transfer the cell type labels from snRNA-seq to snATAC-seq (Methods). Consistently, all major cell types were also present in snATAC-seq (Fig. 1d and Supplementary Fig. 1c). Additionally, we confirmed cell type identities by examining chromatin accessibility at the promoter regions of known markers and calculating a gene activity score that quantified chromatin accessibility within the gene body and promoter regions (Fig. 1e and Supplementary Fig. 1d). Furthermore, we identified cell type-specific differentially expressed genes (DEGs) and differentially accessible chromatin regions (DARs) for each cell type (Supplementary Table 2 and 3), and found that the cell types can be well-distinguished by those DEGs and DARs (Supplementary Figs. 1e,f).

**Fig.1:**
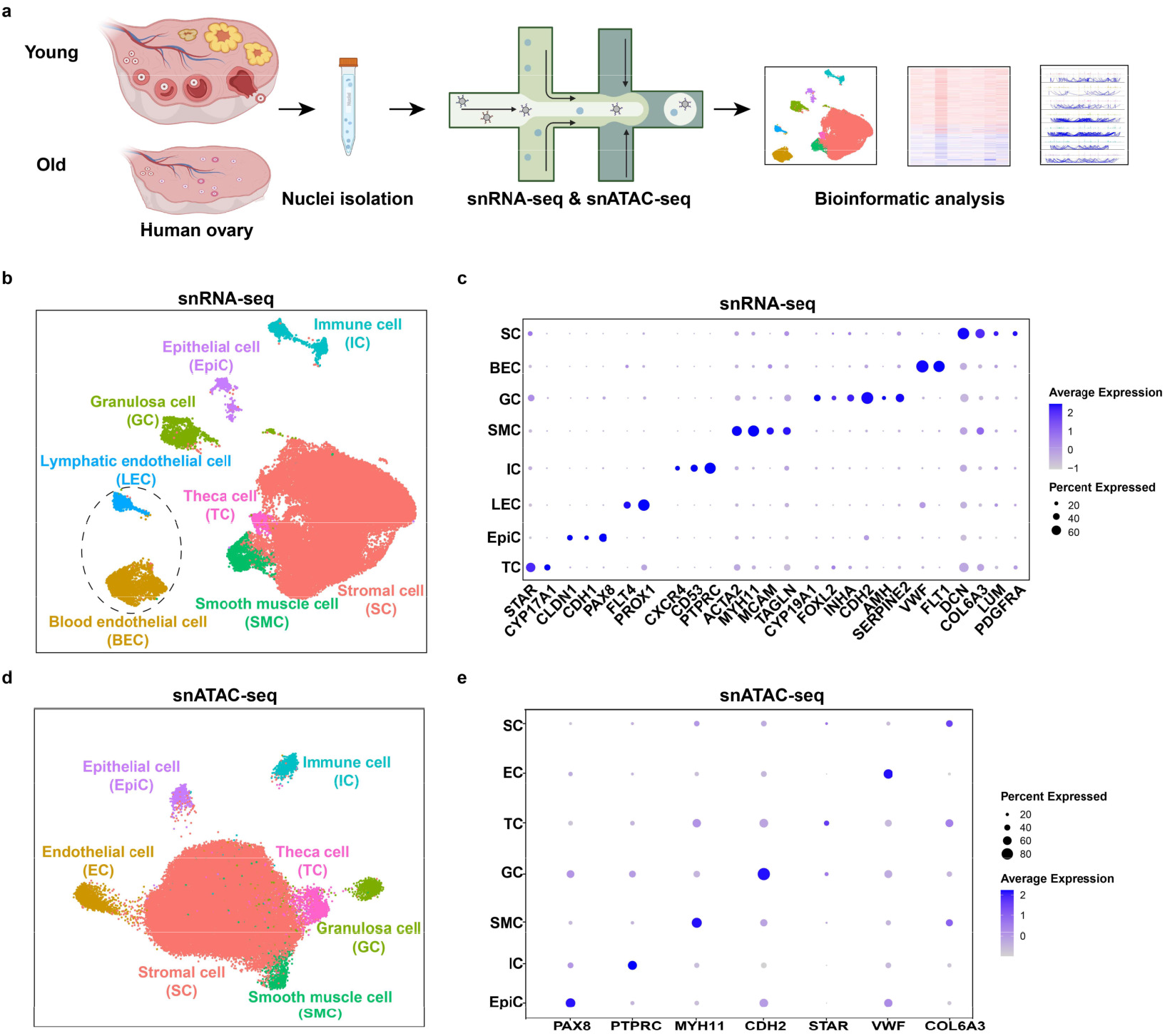
Single-nuclei transcriptomic and chromatin accessibility profiling of the human ovary. **a**, Schematic representation of experimental methodology. **b**, UMAP plots of human ovary snRNA-seq dataset. **c**, Dot plot representing relative mRNA expression of selected known markers for each cell type. Dot size indicates the proportion of cells in the cluster expressing a gene, the shading indicates the relative level of expression (low to high reflected as light to dark). **d**, UMAP plots of human ovary snATAC-seq dataset. **e**, Dot plot representing relative gene activity of selected known markers for each cell type. Dot size indicates the proportion of cells in the cluster expressing a gene, the shading indicates the relative level of expression (low to high reflected as light to dark).

### Altered cell type composition with age

To investigate the dynamic changes in cell type composition during human ovarian ageing, we compared the cell type proportions of aged and young ovaries in the snRNA-seq data. We found significant changes in the proportions of several cell types during ageing (Fig. 2a). For example, the abundance of granulosa and theca cells, two critical components of ovarian follicles, were significantly decreased in aged compared to young ovaries (Fig. 2a), in line with the well-known phenomenon of decreasing follicle number with increasing age^32^. In addition, blood vessel and lymphatic endothelial cells, the cell layers lining the blood and lymph vessels, respectively, also markedly decreased in proportion in aged ovaries (Fig. 2a), consistent with the observed negative correlation between ovarian vascularity and age^33^. Interestingly, epithelial cells were the only cell type that increased in proportion in aged ovaries (Fig. 2a), potentially reflecting the lifetime of ovulation-induced rupture and repair experienced by aged ovaries^34^. Consistently, aged epithelial cells exhibited signicantly elevated expression of cell cycle-associated genes, while aged granulosa cells, theca cells, and endothelial cells exhibited increased expression of apoptosis-associated genes and/or decreased expression of cell cycle-associated genes compared to young couterparts in the ovary (Supplementary Fig. 2a). In agreement with the snRNA-seq results, we observed almost identical age-related changes in the cellular composition estimated from the snATAC-seq data (Fig. 2b). These results indicated that ageing significantly remodels the cellular architecture of the human ovary.

**Fig.2:**
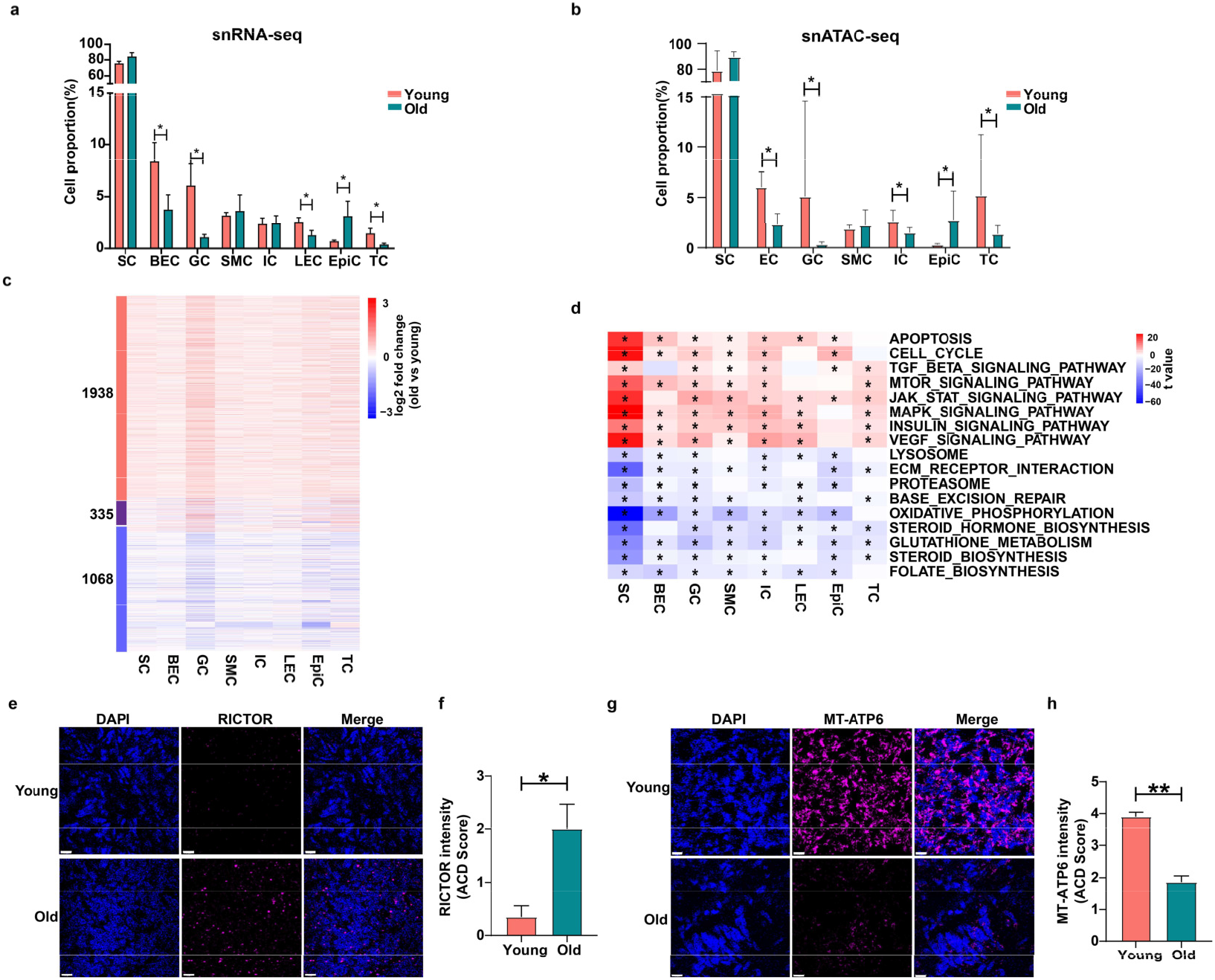
Ageing alters ovarian cellular composition and affects the transcriptional activity of pathways involved in the hallmarks of ageing across cell types. **a**, Bar plots represent the proportion of each cell type in young and aged ovaries estimated from snRNA-seq data. (Mean±SE; Permutation test; Asterisk (*) indicates FDR<0.05 and abs(log2FD)>1.5; Methods). **b**, Bar plots represent the proportion of each cell type in young and aged ovaries estimated from snATAC-seq data. (Mean±SE; Permutation test; Asterisk (*) indicates FDR<0.05 and abs(log2FD)>1.5). **c**, Heat map displaying log2 fold changes in gene expression (aged vs. young) of human ovarian ageing-associated DEGs in each cell type. **d**, Heat map showing selected up- and down-regulated pathways significantly altered in at least 6 cell types during human ovarian ageing. Asterisk (*) indicates a statistically significant difference (*P*_adj_ <0.05). **e**, Representative *in situ* hybridization (RNAscope) images from fresh-frozen human ovary tissue for *RICTOR* staining. **f**, Quantification of *RICTOR* expression in human ovary (young versus old). Mean±SE; n = 2; \**P* < 0.05. **g**, Representative *in situ* hybridization (RNAscope) images from fresh-frozen human ovary tissue for *MT-ATP6* staining. **h**, Quantification of *MT-ATP6* expression in human ovary (young versus old). Mean±SE; n = 2; \*\**P* < 0.01.

### Coordinated changes in ageing hallmarks

To investigate the dynamic changes in gene expression during human ovarian ageing, we identified ageing-associated DEGs for each cell type (Methods). In total, we identified 3,341 ageing-associated DEGs (Supplementary Table 4), the number of which ranged from a few hundred to several thousand, depending on cell type (Supplementary Fig. 2b). Specifically, granulosa cells have the largest number of ageing-associated DEGs (n=2,255) (Supplementary Fig. 2b), suggesting that granulosa cells are more vulnerable to ageing than other cell types in the human ovary. Interestingly, we found that most ageing-associated DEGs are shared among cell types (Supplementary Fig. 2c) and show congruent changes in expression (Fig. 2c). Among the “common DEGs” which were significantly differentially expressed in at least four cell types (Supplementary Fig. 2d), 218 genes were up-regulated and 182 were down-regulated in the aged ovary (Supplementary Fig. 2d). Common DEGs include those reported in the GenAge database as human ageing-related genes, such as *RICTOR, IGF1R, MAP3K5*, and *APOE* (Supplementary Figs. 2e,f). Gene ontology (GO) analysis^35^ indicated that common DEGs were enriched in the “hallmarks of ageing”^36^, including pathways involved in nutrient sensing signaling, cellular senescence, proteostasis, cellular communication, and mitochondrial function (Supplementary Fig. 2g). We also found cell-type-specific ageing-associated DEGs (Supplementary Fig. 2h), such as those enriched in cell type-relevant functions, including vasculogenesis for blood vessel endothelial cells, follicle development for granulosa cells, and smooth muscle contraction for smooth muscle cells (Supplementary Fig. 2i).

To gain insight into the dynamic changes in biological pathways during human ovarian ageing, we used Gene Set Variation Analysis (GSVA)^37^ to estimate the pathway activity score for individual cells and compare the pathway activity between young and aged ovaries in each cell type (Methods). We found that half (92/186) of KEGG pathways were significantly up- or down-regulated in at least six cell types in aged ovaries (Supplementary Fig. 3a). Genes involved in these pathways showed congruent changes in expression direction across cell types (Supplementary Fig. 3a). Notably, expression of genes involved in the nutrient-sensing signaling pathways, including the mTOR, insulin, and MAPK pathways, increased in the aged ovaries across cell types, while those involved in oxidative phosphorylation and base excision repair decreased (Fig. 2d and Supplementary Figs. 3b-e). To validate the age-related changes in gene expression, we performed *in situ* hybridization assays and confirmed the increased expression of the mTOR signaling gene *RICTOR*, and decreased expression of the oxidative phosphorylation gene *MT*-*ATP6* in aged ovaries in vivo (Figs. 2e-h). Together, these results indicate that the human ovary undergoes coordinated transcriptomic changes during ageing, resulting in profound alterations to processes central to the biology of ageing.

**Fig.3:**
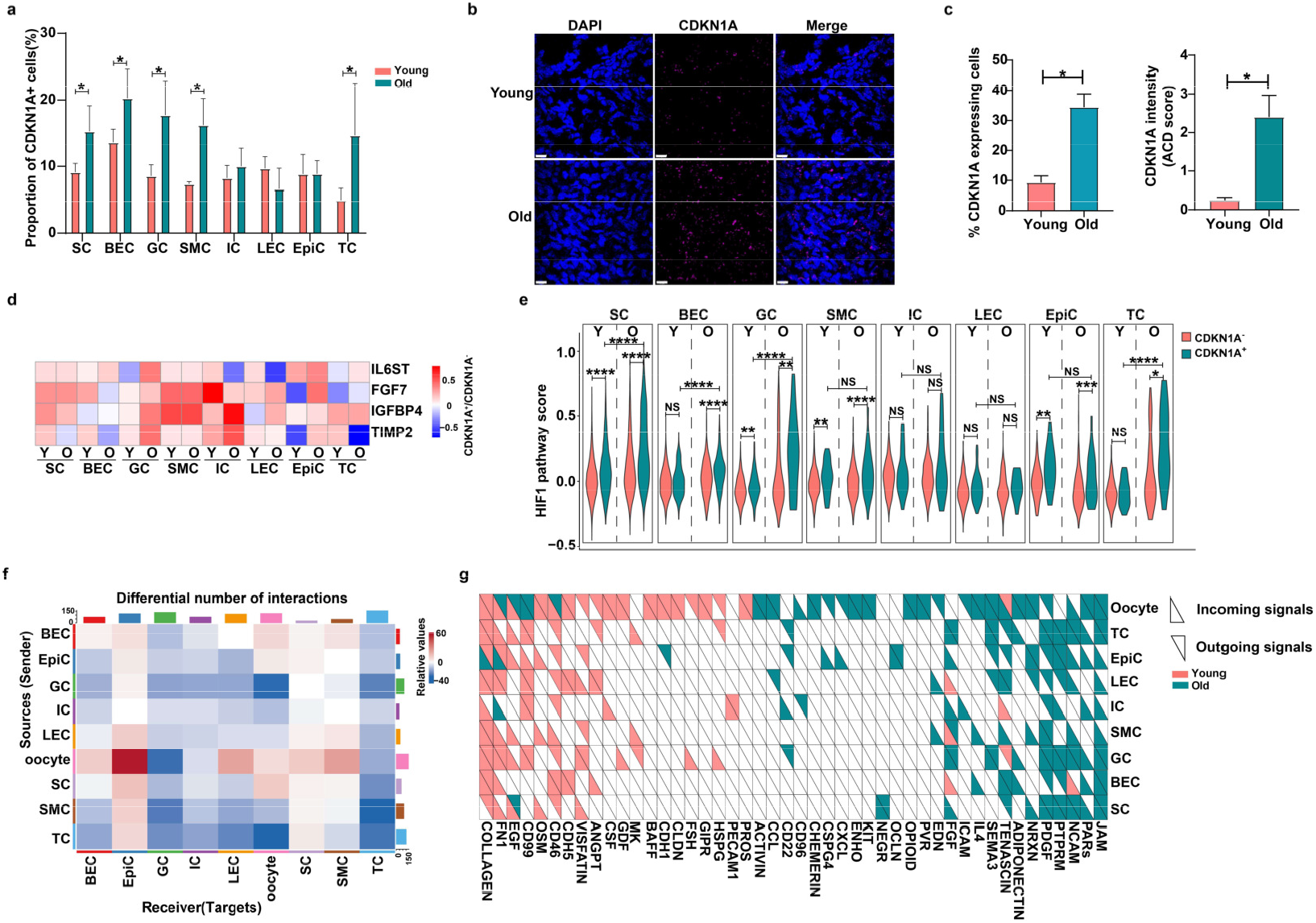
Aging increases signatures of cellular senescence and alters cellular communication in the ovary. **a**, Bar plots represent the proportion of *CDKN1A*^+^ cells for each cell type in young and aged ovaries. (Mean±SE; Permutation test; Asterisk (*) indicates FDR<0.05 and abs(log2FD)>1.5). **b**, Representative *in situ* hybridization (RNAscope) images from fresh-frozen human ovary tissue for *CDKN1A* (p21) staining. **c**, Quantification of *CDKN1A* expression and the proportion of *CDKN1A*^+^ cells in the human ovary (young versus old). Mean±SE; n = 2; \**P* < 0.05. **d**, Heat map displaying log2 fold changes in gene expression (*CDKN1A*^+^ vs. *CDKN1A*^-^ cells) of selected SASP genes in each cell type. **e**, Violin plots showing the module score of HIF-1 pathway genes in *CDKN1A*^+^ cells and *CDKN1A*^-^ cells from each type. (Two-sided Wilcoxon test; NS: Not significant, \**P*<0.05, \*\**P*<0.01, \*\*\**P*<0.001, \*\*\*\**P*<0.0001). **f**, Heat map of the differential number of interactions between cell types in young and aged ovaries. The top bar plots represent the sum of each column of values displayed in the heatmap (incoming signaling). The right bar plots represent the sum of each row of values (outgoing signaling). **g**, Heat map showing the outgoing and incoming signaling pathways significantly enriched in young or aged ovaries for each cell type.

### Cellular senescence in the human ovary

Senescent cell burden increases with age in various tissues in the context of physiological ageing and ageing-related disease^38-41^. To test if cellular senescence increases during human ovarian ageing, we examined the expression of the widely used senescence markers, *CDKN1A* (p21) and *CDKN2A* (p16), in the human ovary. On average, very few (∼0.43%) ovarian cells expressed *CDKN2A*, while a considerable proportion of young ovarian cells expressed *CDKN1A* (∼9.49%) and a significantly higher proportion of *CDKN1A*^*+*^ cells (∼15.56%) was observed in aged ovaries (Supplementary Figs. 4a,b). We then calculated the proportion of *CDKN1A*^*+*^ cells for each cell type in young and aged ovaries. We found a significant increase in the proportion of *CDKN1A*^*+*^ cells with age in stromal, granulosa, theca, blood vessel endothelial, and smooth muscle cells (Fig. 3a). Using *in situ* hybridization, we found a ∼3-fold increase in both the proportion of cells expressing *CDKN1A*, and in the average expression of *CDKN1A*, in aged compared to young ovaries (Figs. 3b,c). In addition, a subset of senescence-associated secretory phenotype (SASP) genes were up-regulated in *CDKN1A*^*+*^ cells in the human ovary (Fig. 3d). To gain insight into the transcriptional signatures of *CDKN1A*^*+*^ cells, we identified the DEGs in *CDKN1A*^*+*^ cells compared to *CDKN1A*^*-*^ cells from both young and aged stromal cells (Supplementary Fig. 4c). GO analysis indicated that genes involved in “response to oxygen level” and “HIF-1 signaling pathway” are up-regulated in *CDKN1A*^*+*^ stromal cells (Supplementary Fig. 4c). We then compared the transcriptomes of *CDKN1A*^*+*^ cells between young and aged stromal cells and found that genes involved in the HIF-1 pathway as well as those in the nutrient-sensing signaling were up-regulated in aged-compared to young *CDKN1A*^*+*^ stromal cells (Supplementary Fig. 4c). To test whether the up-regulation of the HIF-1 pathway is a senescence signature in the ovary, we computed HIF-1 pathway scores based on the HIF-1 pathway-related DEGs that were up-regulated in *CDKN1A*^*+*^ cells (Supplementary Table 5), including the key HIF-1 target genes that regulate NAD^+^ metabolism (*NAMPT*)^42^, cellular respiration (*PDK1*)^43^, and apoptosis (*DDIT4*)^44^ in response to hypoxia (Supplementary Fig. 4d). We found that the HIF-1 pathway was significantly enhanced in *CDKN1A*^*+*^ cells and further elevated during ageing in most cell types (Fig. 3e). Given the reduced vasculature of aged ovaries^33^, our results suggested that the hypoxic environment might be a critical factor in driving cellular senescence in human ovaries, through upregulation of HIF-1 signaling.

**Fig.4:**
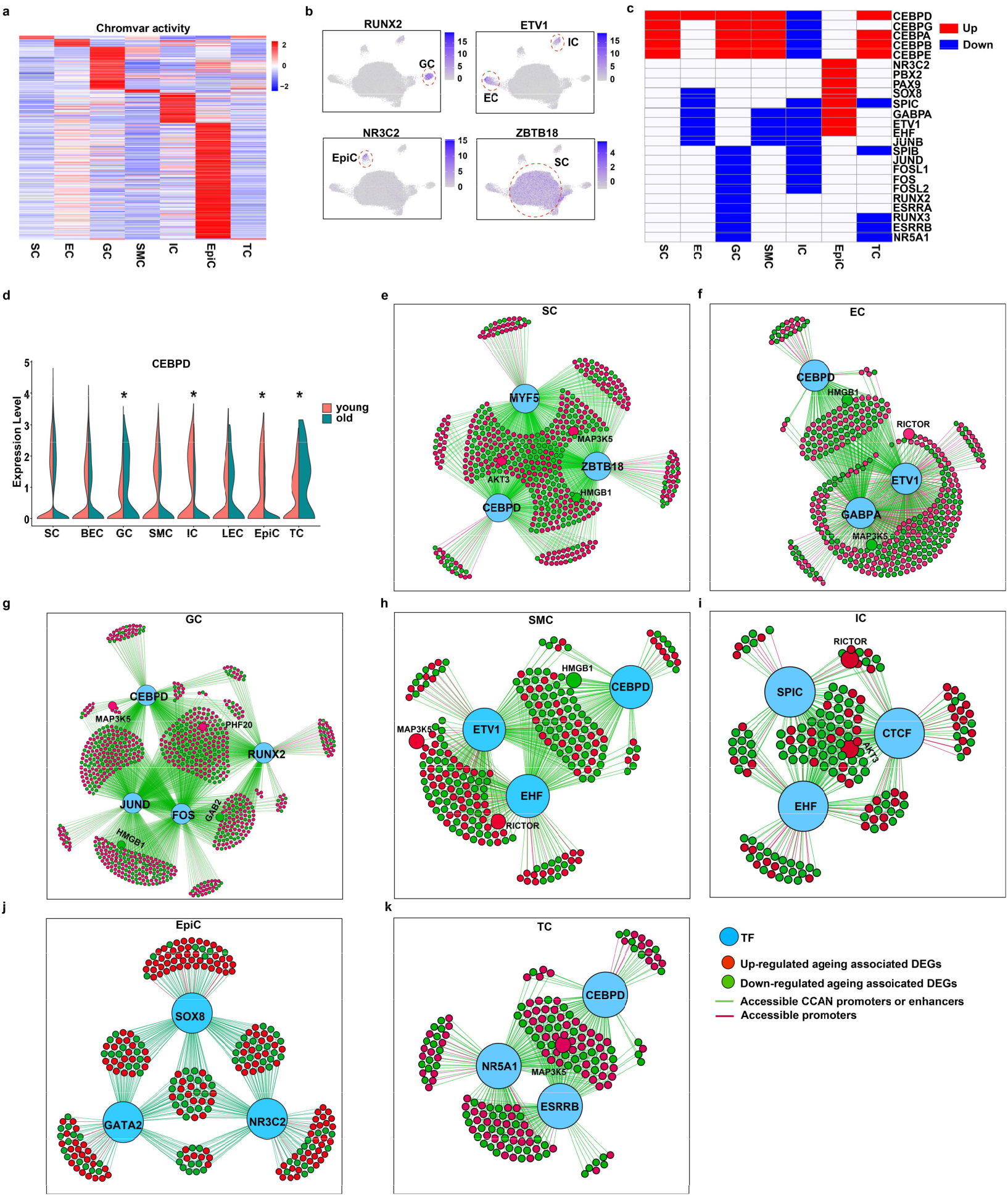
Cell type-specific TF regulatory networks implicate CEBPD as an important regulator of ageing-associated gene expression in the human ovary. **a**, Heat map showing the average chromVAR motif activity for each cell type. **b**, UMAP plots displaying the chromVAR motif activity of selected cell type-specific TFs. **c**, Heat map showing the TFs with significant changes in chromVAR motif activity in each cell type during ovarian ageing. **d**, Split violin plots showing the expression levels of *CEBPD* in each cell type from young and aged ovaries. (MAST test; \**P*_adj_<0.05). **e-j**, TF regulatory network plots showing the top regulators of ageing-associated DEGs in each cell type.

### Ageing alters cellular communication

Altered intercellular communication is a hallmark of ageing^36^. To explore potential age-related alterations to the ovarian cellular communication network, we used CellChat^45^, which models the probability of the cell-cell interaction network based on gene expression and prior knowledge of ligand-receptor interactions. To gain insight into the intercellular communication between ovarian somatic cells and oocytes, we integrated our snRNA-seq with publicly available human oocyte single-cell transcriptomes from young and reproductively aged ovaries^46^. We found that ageing slightly reduced both the total number and overall strength of intercellular interactions (Supplementary Fig. 5a,b). Strikingly, aged granulosa and theca cells received far fewer signals from all other cell types, and the same trend was observed for blood/lymphatic endothelial cells and immune cells (Fig. 3f). Of particular relevance to fertility, the number of signals received by oocytes from granulosa and theca cells profoundly decreased with age (Fig. 3f), consistent with evidence that the functions of granulosa and theca cells in supporting oocytes fail during ageing^47^. In contrast, the interaction number and strength from all cell types to epithelial cells increased (Fig. 3f and Supplementary Fig. 5c). We further identified the incoming and outgoing signaling pathways that exhibited a significant difference in communication probability between young and aged ovaries for each cell type (Fig. 3g). In total, we identified 46 pathways with significant differential communication probability with age (Fig. 3g). Notably, the COLLAGEN and FN1 (Fibronectin) pathways, core components of extracellular matrix (ECM) biology, exhibited a significantly higher communication probability in most cell types in the young ovary, suggesting an essential role of the ECM in maintaining ovary function. Interestingly, epithelial cells were the only cell type that exhibited a significantly higher communication probability of COLLAGEN and FN1 signaling in the aged ovary (Fig. 3g). COLLAGEN and FN1 signaling is known to promote the proliferation of epithelial cells^48,49^, suggesting that activation of these pathways may explain the increased proportion of epithelial cells we observed in the aged ovary (Figs. 2a,b). In contrast, the JAM (Junction adhesion molecule), PARs (Protease-activated receptors), and NCAM (Neural cell adhesion molecule) pathways, which mediate cell-cell adhesion processes, exhibited a significantly higher communication probability in most cell types in the aged ovary (Fig. 3g). The PDGF pathway, which has known roles in fibroinflammatory processes, was significantly enriched in all the cell types in the aged ovary (Fig. 3g), consistent with the finding of elevated fibroinflammatory cytokines in aged human ovarian follicular fluid^50^. In contrast to the age-related loss of interactions between granulosa and theca cells and oocytes (Fig. 3f), several known signaling pathways that are critical to maintaining the function of oocytes and granulosa cells were specifically enriched in young oocytes and granulosa cells, including the FSH and GDF pathways that maintain follicle growth and function^51^ (Fig. 3g). Our results demonstrate that human ovarian ageing is characterized by significant changes in cellular communications among oocytes and somatic cell types, potentially contributing to an age-related loss of follicular function, tissue fibrosis, and epithelial hyperplasia.

**Fig.5:**
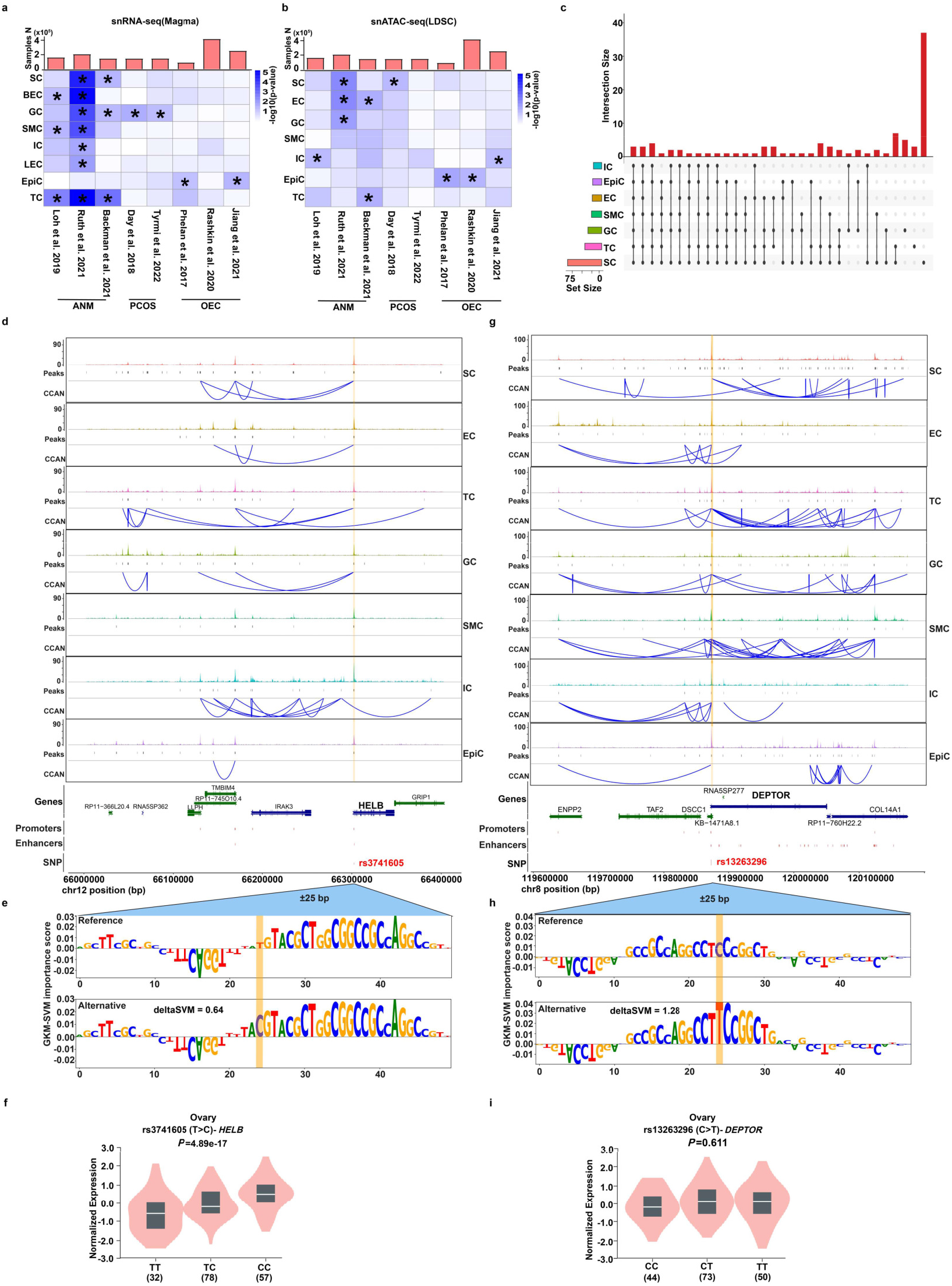
Integration of ANM GWAS, single-nuclei multi-omics, and machine-learning nominates causal variants and gene targets associated with human ovarian aging. **a**, Heat map of enrichment significance of ovary-relevant trait GWAS variants in ovary cell type gene expression signatures. **b**, Heat map of enrichment significance of ovary-relevant trait GWAS variants in ovary cell type-specific chromatin accessibility. **c**, Upset plot showing the intersection size between sets of ANM-associated SNPs that overlap with transcriptional regulatory elements found in each cell type. The bar plot on the left shows the set size of variants for each cell type, and the bar plot on the top shows the number of overlapping SNPs shared by two or more sets, or the number of unique variants in one set. **d**, Cis-regulatory architecture at the *HELB* gene in each cell type. The snATAC-seq tracks represent the aggregate signals of all cells from a given cell type. The co-accessible peaks inferred by Cicero for each cell type are shown. **e**, The gkm-SVM importance score for each base within the ±25-base pair (bp) region surrounding rs3741605. **f**, The eQTL effect of rs3741605 on *HELB* expression in human ovary tissue from the GTEx database. **g**, Cis-regulatory architecture at the *DEPTOR* gene in each cell type. The snATAC-seq tracks represent the aggregate signals of all cells from a given cell type. The co-accessible peaks inferred by Cicero for each cell type are shown. **h**, The gkm-SVM importance score for each base within the ±25-base pair (bp) region surrounding rs13263296. **i**, The eQTL effect of rs13263296 on *DEPTOR* expression in human ovary tissue from the GTEx database.

### Ageing alters cell identity TF networks

Master transcription factors (TFs) largely determine cell identity^52^. As loss of cell identity with age has been implicated in age-related tissue dysfunction, we next identified cell identity-associated TFs, and investigated for age-related changes in their activity (Methods) in the snATAC-seq data. As expected, individual cell types can be distinguished by predicted motif activity (Fig. 4a and Supplementary Table 6). The motifs of folliculogenesis-related TFs, mainly AP-1 and RUNX transcription factors^53^, were predominantly enriched in granulosa cells (Fig. 4b and Supplementary Fig. 6a). Steroidogenesis-related TFs^54,55^ were mainly enriched in granulosa cells and theca cells (Supplementary Fig. 6b). The TF footprinting analysis highlighted cell type-specific enrichment of those TFs in granulosa cells and theca cells (Supplementary Fig. 6c). ETS TFs are central regulators of endothelial and immune cells^56,57^, both of which originate from the hemogenic endothelium during embryogenesis^58^. Consistently, a family of ETS TFs were enriched in endothelial and immune cells (Fig. 4b and Supplementary Fig. 6d). Additionally, we identified the TFs enriched in epithelial cells and stromal cells, respectively (Fig. 4b and Supplementary Fig. 6e).

To reveal the TFs that govern human ovarian ageing, we compared the predicted motif activity between young and aged cells in each cell type (Supplementary Table 7). Surprisingly, CCAAT/enhancer binding proteins (C/EBPs) motif activities significantly increased in most cell types except immune cells and epithelial cells (Fig. 4c). Among the members of this TF family, *CEBPD* was highly expressed across cell types, and its age-related changes in expression were in line with the changes in motif activity during ovarian ageing (Fig. 4d). In addition, most cell identity-associated TFs exhibited significantly decreased motif activity, while epithelial cell identity-associated TFs exhibited significantly enhanced motif activity during ovarian ageing (Fig. 4c). We further calculated cell identity scores in each cell type by examining the expression level of the top 100 cell type-specific genes (Methods). We found that cells in young ovaries exhibited high expression of their corresponding cell type-specific genes and minimal expression of other cell type-specific genes (Supplementary Fig. 6f). Surprisingly, granulosa cells, immune cells, and theca cells in aged ovaries expressed deficient levels of their corresponding cell type-specific genes, and instead expressed high levels of stromal cell-specific genes (Supplementary Fig. 6f). These results suggest a prevalent loss of cell identity in aged ovaries.

We next sought to build cell type-specific TF regulatory networks for human ovarian ageing, and first constructed the cis co-accessibility networks (CCANs) in each cell type using Cicero^59^. Next, we defined the putative enhancers and promoters by overlaying the CCAN peaks with human ovary tissue enhancer and promoter annotations from the ENCODE database^60^. Finally, we defined a gene as a CCAN-linked gene if one of the CCAN peaks lies in its putative promoter (Supplementary Fig. 7a). In this way, we identified varying numbers of CCANs and CCAN-linked genes for each cell type (Supplementary Fig. 7b). Most cell type-associated DEGs and ageing-associated DEGs significantly overlapped with CCAN-linked genes in all cell types (Supplementary Fig. 7c), suggesting that CCANs play essential roles in determining cell identity and the regulation of ovarian ageing. For each cell type, we built the ageing-associated TF regulatory networks governed by the top TFs (n=3∼4) that change with age, as defined by the predicted motif activity. Those ageing-associated DEGs whose promoters or putative enhancers contained both accessible peaks and motifs of the top ageing-associated TFs within the peaks were defined as candidate targets of the selected TFs (Supplementary Fig. 7a). We generated the ovarian ageing-associated TF regulatory network for each cell type and found a critical role for CEBPD in human ovarian ageing (Figs. 4e-k). We found that CEBPD target genes are enriched in processes of known importance to the basic biology of ageing, including mTOR signaling, MAPK signaling, and cellular senescence, in multiple cell types (Supplementary Fig. 7d).

### Cellular targets of ANM genetic risk

The most comprehensive recent ANM GWAS identified 290 ANM-associated genetic risk loci^16^. Using gene expression data from several publicly-available datasets, the Ruth et al. study implicated hematopoietic stem and progenitor cells as the major cellular targets of ANM-associated risk variants^16^. Since the Ruth et al. analysis did not include single-cell data from the human ovary, we next investigated whether any specific cell types in the human ovary were enriched for ANM-associated variants, by performing MAGMA^61,62^ analysis using our snRNA-seq dataset. For comparison, we also included GWAS data from 2 other ovarian phenotypes, ovarian epithelial cancer (OEC) and polycystic ovary syndrome (PCOS). We found that ANM-associated variants were significantly enriched in almost all cell types (Fig. 5a), indicating a systemic effect of ageing on the ovary. In contrast, OEC- and PCOS-associated variants were enriched in epithelial cells and granulosa cells, respectively (Fig. 5a). To investigate if any cell type-specific regions of chromatin accessibility were enriched for ANM-associated variants, we also performed cell type-specific linkage disequilibrium (LD) score regression^63^ using our snATAC-seq dataset. Consistently, ANM-associated variants showed a significant enrichment in multiple cell types (Fig. 5b). Together with the coordinated changes in transcriptomes and chromatin accessibility we observed across cell types, the results from our analyses of ANM genetic signal suggest that all ovarian cell types contribute to ovarian ageing.

### Functional ANM variants and genes nominated

To gain insights into how ANM-associated variants contribute to ovarian ageing, we performed post-GWAS analyses to identify functional regulatory variants and affected target genes. Using 290 ANM-associated GWAS lead variants from Ruth et al.^16^, we first compiled a comprehensive set of coinherited variants based on LD (R^2^ value ≥0.8) calculated from phase 1 genotypes of individuals of European ancestry in the 1000 Genomes dataset (Methods). In total, we identified 5,555 ANM-associated variants (Supplementary Table 8). To identify the functional variants that may affect transcriptional regulatory activity in each cell type, we first overlapped ANM-associated variants with the putative enhancers and promoters we identified in each cell type. In this way, we identified 101 candidate functional variants (Supplementary Fig. 8 and Supplementary Table 9) and found that a substantial number of these variants were shared across several cell types (Fig. 5c). Next, we focused on the putative functional variants that were shared in at least four cell types, among which 6 variants occured in DNA damage response (DDR)-related gene loci (Fig. 5d and Supplementary Figs. 9a-e). The DDR is the major pathway linked to ovarian ageing as detected by GWAS of ANM^16^ and our results highlight the functional role of ANM-associated variants on the regulation of DDR across major ovarian cell types. For example, the rs3741605 (T>C) allele, associated with delayed ANM (BETA>0; Supplementary Table 10), occurs in the putative promoter of the *HELB* gene, encoding DNA helicase B^64^, that is active in most cell types in the ovary, including stromal, endothelial, theca, granulosa, and immune cells (Fig. 5d). Previously, multiple missense variants predicted to be deleterious have been identified in *HELB* that are associated with early ANM^16^. Remarkably, expression quantitative trait loci (eQTL) analysis from the GTEx database^65^ indicated that the C allele of rs3741605 was significantly correlated with increased expression of *HELB* in the human ovary (Fig. 5f). This result suggest that the functional non-coding variant rs3741605 may contribute to delayed ANM by upregulating the expression of a critical DNA repair gene, thereby conferring improved genome maintenance.

To explore the potential mechanisms underlying the influence of variants on gene expression, we predicted the effect of candidate functional variants on TF binding activity by applying gapped k-mer support vector machine-based methods (LS-GKM^66^ and deltaSVM^67^) (Methods). We found that the delayed ANM-associated C allele of rs3741605 could enhance TF binding activity (Fig. 5e), showing a high concordance of predicted beneficial allelic effect on the increased expression of *HELB* and delayed ANM through enhanced genome maintenance.

We also identified functional regulatory variants with predicted deleterious effects on ANM. The variant rs13263296 is located in the *DEPTOR* locus, a key gene involved in mTOR signaling (Fig. 5g and Supplementary Fig. 10a), and the T allele of rs13263296 is associated with early ANM (BETA<0; Supplementary Table 10) and occurs in the putative transcription start site (TSS)-proximal enhancer of the *DEPTOR* gene in most cell types (Fig. 5g). We observed significant up-regulation of *DEPTOR* expression in aged granulosa cells and epithelial cells (Supplementary Fig. 10b). Of note, rs13263296 (C>T) was not significantly associated with increased *DEPTOR* expression in the human ovary (Fig. 5i), perhaps due to the use of bulk tissue in the eQTL analysis. Mechanistically, the deltaSVM analysis suggested that rs13263296 (C>T) may affect *DEPTOR* expression by enhancing TF binding activity (Fig. 5h). In addition to these loci, we identified several functional regulatory variants located in oxidative phosphorylation and MAPK signaling-related gene loci (Supplementary Figs. 10c,d). Taken together, our integrated analyses revealed the global effects of ANM-associated non-coding variants on gene expression across ovarian cell types and nominated functional regulatory ANM risk variants that may dysregulate genes involved in pathways of relevance to the canonical hallmarks of ageing, such as mTOR and genome maintenance.

In summary, our single nuclei multi-omic analysis of young and reproductively aged ovaries provides high-resolution characterization of the transcriptional regulatory landscape at the single-cell level, uncovering conserved mechanisms of ageing biology in action, such as the hyperactivity of mTOR observed across all ovarian somatic cell types with age. Our results raise the hope that geroprotectors targeting the basic biology of ageing, such as mTOR signaling, may be used to delay reproductive ageing in women. Furthermore, our integrative post-GWAS analyses of ANM provides biological insights into the role of inherited non-coding variants in ovarian ageing in humans, nominating new functional variants for follow-up interrogation. These findings expand our understanding of inherited variation in ANM and provide a roadmap for the functional dissection of the non-coding genetic variation influencing ANM, pointing towards the nomination of new therapeutic targets for reproductive health in women.

## Methods

### Sample procurement

Fresh-frozen healthy human ovary samples were purchased from BioIVT (Baltimore, MD) and Cureline (Brisbane, CA). All samples were de-identified and the tissue source was anonymous to the researcher.

### Nuclear dissociation and library preparation

For snATAC-seq, nuclei isolation was performed according to the 10× Genomics protocol CG000212 (Rev B) with ∼100 mg frozen human ovary sample. Libraries were generated by using 10x Chromium Single Cell ATAC Reagent Kits (v1) and sequenced using the Illumina NextSeq 550 platform with 150-bp paired-end sequencing.

For snRNA-seq, the nuclei isolation was performed according to 10× Genomics protocol CG000393 (Rev A) with ∼100 mg frozen human ovary sample. Libraries were generated using 10x Chromium Single Cell 3’ Reagent Kits (v3) and sequenced using the Illumina NextSeq 550 platform with 150-bp paired-end sequencing.

### Processing of snRNA-seq data

Reads were aligned to a pre-mRNA GTF built on the GRCh38 genome using Cellranger (v3.1.0) to account for unspliced nuclear transcripts. The Cellranger function aggr was used to aggregate all snRNA-seq libraries without depth normalization to generate a gene by nucleus matrix. Nuclei with fewer than 200 genes, nuclei with more than 6000 genes, or nuclei with more than 15% of unique molecular identifiers stemming from mitochondrial genes were removed. In total, we obtained 42,568 nuclei for downstream analysis. Expression levels were normalized with the LogNormalize method in Seurat^28^ (v4.0.4), and the top 2100 highly variable genes (HVGs) were used for principal component analysis (PCA). To remove batch effects, the first 15 PCs were batch corrected using Harmony^29^ (v0.1). Clustering was performed by constructing a K-nearest neighbor (KNN) graph with corrected PCs and applying the Louvain algorithm. Dimensional reduction was performed with Uniform Manifold Approximation and Projection (UMAP) and individual clusters were annotated based on expression of cell type-specific markers. Differentially expressed genes (DEGs) were identified with the Seurat FindMarkers function for genes detected in at least 25% of cells, using the MAST test and a log-fold-change threshold of 0.25. Bonferroni-adjusted p-values were used to determine significance at *P*_adj_ < 0.05. Gene ontology analysis was performed using Metascape^35^. Cell cycle score, apoptosis score, HIF pathway score, and cell identity score were evaluated by AddModuleScore function in Seurat with corresponding gene lists, respectively.

### Processing of snATAC-seq data

Reads were aligned to the GRCh38 genome using cellranger-atac (v2.0.0) Libraries were aggregated with cellranger-atac without depth normalization to generate a peak by nucleus matrix. Low-quality nuclei (peak region fragments < 200, peak region fragments > 10000, percentage of reads in peaks > 8, blacklist ratio < 0.01, TSS enrichment > 1.5 & nucleosome signal < 1.5) were removed using Signac^30^ (v1.5.0). In total, we obtained 41,550 nuclei for downstream analysis. The peak by cell matrix was transformed using the term frequency-inverse document frequency (TF-IDF). Dimensional reduction was performed via singular value decomposition (SVD) of the TF-IDF matrix. The first 40 latent semantic indexing (LSI) components were batch corrected using Harmony (v0.1). Clustering was performed by constructing a K-nearest neighbor (KNN) graph with corrected LSI components and applying the Louvain algorithm. Peak calling was performed with the CallPeaks function in MACS2^68^ (v2.2.7.1) in each cluster. Gene activity was estimated by counting ATAC peaks within the gene body and 2 kb upstream of the TSS using protein-coding genes annotated in the Ensembl database. Canonical correlation analysis^31^ (CCA) was used to capture the shared feature correlation structure between snATAC-seq gene activity and snRNA-seq gene expression. Mutual nearest neighbors^31^ (MNNs) were then identified the pairs of corresponding cells that anchor the two datasets together. We assigned the cell types to the snATAC-seq clusters if the majority (>80%) of cells were aligned to the corresponding cell type. Differentially accessible chromatin regions (DARs) between cell types were assessed with the FindMarkers function for peaks detected in at least 5% of cells, using the MASTtest and a log-fold-change threshold of 0.25. Bonferroni-adjusted p-values were used to determine significance at P_adj_ < 0.05. The single-nuclei motif activity for a set of 452 human TFs from the JASPAR 2020^69^ was computed by running chromVAR (v1.14.0) through the RunChromVAR function in Signac. Differential motif activity between young and old ovaries in each cell type was identified by using the FindMarker function for chromVAR motifs detected in at least 25% of cells, using the MAST test and a log-fold-change threshold of 0.50. Bonferroni-adjusted p-values were used to determine significance at Padj < 0.05. To further analyze specific TFs of interest, we used the Footprint function in Signac to perform TF footprinting analysis.

### Gene set variation analysis (GSVA)

Pathway analyses were performed on the 186 Kyoto Encyclopedia of Genes and Genomes (KEGG) pathways in the MSigDB^70^ database (v7.4.1). GSVA^37^ (v1.40.1) was used to perform gene set variation analysis to estimate the pathway activity score for individual cells. To compare the pathway activity scores between young and old ovaries in each cell type, we contrasted the activity scores using the limma^71^ package (v3.48.3). Bonferroni-adjusted p-values were used to determine significance at an P_adj_ < 0.05. T-values of pathways that exhibited significance in at least 6 cell types were visualized using heatmaps.

### Generation of cis-coaccessibility networks with Cicero

We applied Cicero^59^ (v1.3.4.11) to generate cis-accessibility networks (CCANs) for each cell type. Briefly, the Signac object for each cell type was converted to the CellDataSet format and then made into a Cicero object. The algorithm assigned the cells into many groups, each group comprised of 50 cells similarly positioned in clustering space. Graphical LASSO was used to calculate the correlations in adjusted accessibilities between all pairs of ATAC peaks within 500 kb. Finally, CCANs were identified through community detection.

### Transcription factor regulatory network construction

For a given TF, the ovarian ageing-associated DEGs whose promoters or putative enhancers contained both accessible peaks and motifs of the certain ageing-associated TFs within the peaks were defined as candidate targets of the selected TFs. We used this information to construct a directed TF regulatory network using the Gephi (v0.9.2).

### Cell type enrichment analysis

For the snRNA-seq data, to estimate the association of gene-level GWAS trait association statistics with gene expression specificity in a given cell type, we used EWCE^72^ (v1.0.0) to calculate gene expression specificity in each cell type. Then, MAGMA.Celltyping^62^ (v1.0.0) was used to calculate the quantile groups for each cell type with the prepare.quantile.groups function. The GWAS variants were then annotated onto their neighbouring genes (Genes were extended 10 kb upstream and 1.5 kb downstream). Finally, MAGMA^61^ (v1.08) was used to test for a positive association (one-sided test) between the cell type specificity and the gene-level associations. *P*-values were used to determine significance at *P* < 0.05.

For the snATAC-seq data, we used ldsc^63^ (v1.0.1) to annotate each variant according to whether or not it overlapped ATAC peaks in each cell type for each GWAS summary statistics. We then estimated partitioned LD scores with the annotated files, HapMap SNPs, and PLINK data corresponding to 1000 genomes phase 3. The baseline model was downloaded from ldsc website. Finally, we used stratified LD score regression to assess the contribution of an annotation to each GWAS trait heritability. *P*-values were used to determine significance at *P* < 0.05.

All GWAS summary statistics for age at menopause^16,73,74^, polycystic ovary syndrome (PCOS)^75,76^, and ovarian epithelial cancer (OEC)^77-79^ were downloaded from GWAS Catalog (https://www.ebi.ac.uk/gwas/) or ReproGen (https://www.reprogen.org/), and re-formatted with MungeSumstats (v1.3.5) or munge_sumstats.py in the ldsc package.

### The effect of variants on transcription factor binding activity

To predict the TFs binding activity score, we overlapped our ovary ATAC peaks with human ovary tissue enhancer and promoter annotations from the ENCODE database. In this way, we obtained 71,470 putative enhancers and promoter regions, which were used as the positive set. We generated the random length and GC-matched genome sequences as the negative set. We then used the gkmtrain function from LS-GKM (v0.1.1)^66^, a new gkm-SVM software for large-scale datasets, to train the TFs binding model for human ovary with positive set, negative set, and “gkmrbf” kernel. For the variants of interest, we retrieved the ±25 bp reference DNA sequence around the variant. To generate the corresponding alternative DNA sequence, we replaced the 25th position with the effect allele. To compute deltaSVM scores, we generated all possible non-redundant k-mers of size 11 and scored each of them using the trained model. We then used deltaSVM to compute the deltaSVM scores with k-mer scores, reference sequences, and alternative sequences. For the GkmExplain scores, we used GkmExplain^80^ on the reference sequences or alternative sequences of variants of interest. The GkmExplain scores were visualized using logomaker (v0.8) (https://github.com/jbkinney/logomaker).

### Cellular communication

To build cell-cell interactions between somatic cells and oocytes in young and old human ovaries, we used CellChat^45^ (v1.1.3) to infer the cell-cell interactions based on the expression of known ligand-receptor pairs in different cell types with a combination of our snRNA-seq datasets and publicly available human oocyte single-cell RNA-seq datasets^46^ from reproductive young and old females. Briefly, we inferred the cell-cell interactions for young and old ovaries, respectively. Next, we used “rankNet” function in CellChat to identify the significant outgoing or incoming signaling enriched in young or old ovaries.

### *In situ* hybridization assay

Flash-frozen human ovary tissues were sectioned at 10 μm. RNA in situ hybridization was performed using RNAscope Multiplex Fluorescent v2 kits (Advanced Cell Diagnostics) according to the manufacturer’s instructions, except fixed with 4% PFA 90 mins at RT and protease IV incubation was performed for 15 min. Probes used were MT-ATP6 (532961), RICTOR (544841), and CDKN1A (311401). Fluorophores used were Opal 690 (1:1500 dilution, Perkin Elmer). Images were taken on a Leica TCS SP8 MP at ×40 magnification. 4-10 regions per sample were analyzed using HALO (v3.2). The mRNA expression levels were evaluated according to the ACD scoring system (https://acdbio.com/dataanalysisguide) by counting number of dots per cell. The cells with at least 4 dots were recognized as *CDKN1A*^+^ cells.

### Reporting Summary

Further information on research design is available in the Nature Research Reporting Summary linked to this article.

## Data availability

The snRNA-seq and snATAC-seq data reported in this paper have been deposited in the Gene Expression Omnibus (GEO) under accession numbers: GSE202601. All other data supporting the findings of this study are available from the corresponding authors on reasonable request.

## Code availability

The codes used to analyze the snRNA-seq and snATAC-seq data are available at https://github.com/ChenJin2020/The-regulatory-landscapes-of-human-ovarian-ageing.

## Acknowledgments

We thank Wilber Quispe (SingulOmics Corporation) for snRNA-seq and snATAC-seq libraries preparation and Columbia Genome Center for sequencing. This work was supported by NIH grants AG069750, DK127778, AG057433, AG061521, HL150521, AG055501, AG057341, AG057433, AG057706, AG057909, and AG17242 (Y.S), a grant GCRLE-1320 (Y. S.) from the Global Consortium for Reproductive Longevity and Equality at the Buck Institute, made possible by the Bia-Echo Foundation, and a grant from The Simons Foundation (Y.S.).

## Author information

These authors contributed equally: Chen Jin, Xizhe Wang

## Contributions

C.J. designed and performed the experiments, analyzed the data, and wrote the manuscript; X.W. performed the *in situ* hybridization assay, and analyzed the imaging data; A.H. provided the analytic advice and revised the manuscript; A.G. generated part of snRNA-seq libraries; M.P. assisted the snRNA-seq data analysis; S.K. identified the LD SNPs and provided the analytic advises; D.C. assisted the *in situ* hybridization assay; J.H. and J.C. revised the manuscript; R.L, Z.W. and M.R. provided conceptual advice; Y.S. conceived and designed the research, analyzed and interpreted the data, and wrote the manuscript.

## Corresponding author

Correspondence to Yousin Suh

## Ethics declarations Competing interests

The authors declare no competing interests

