## Supplemental Figures for "The regulatory landscapes of human ovarian ageing"

**Supplementary Fig.1**

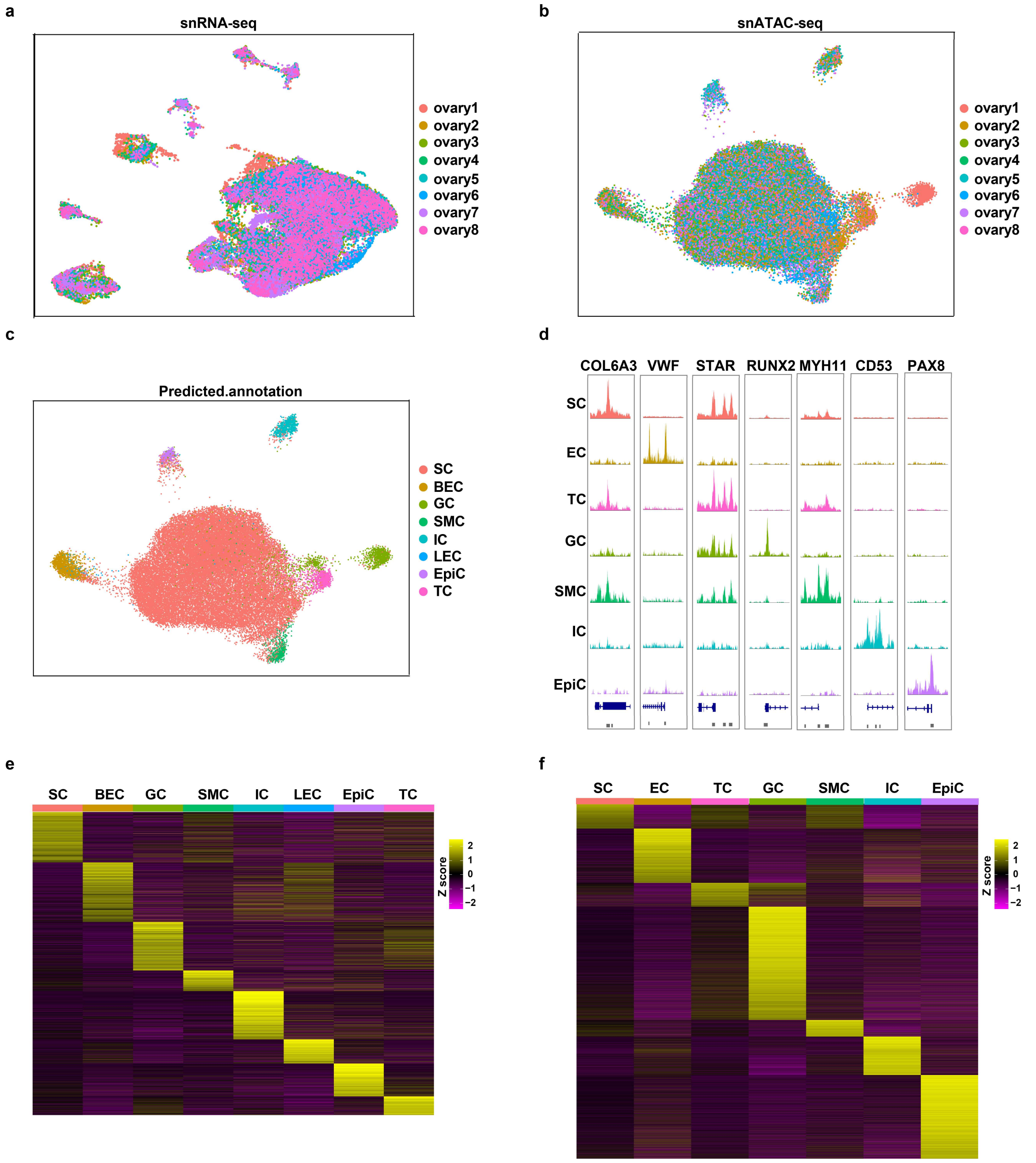


### Supplementary Fig.1: Cell type assignments for human ovary snRNA-seq and snATAC-seq datasets

**a,** UMAP plots of human ovary snRNA-seq dataset. Cell colors based on the donors are shown. **b**, UMAP plots of human ovary snATAC-seq dataset. Cell colors based on the donors are shown. **c,** UMAP plots of human ovary snATAC-seq dataset. Cell colors based on the snRNA-seq predicted annotation are shown. **d,** Genome browser plots showing the pseudo-bulk chromatin accessibility profiles for each cell type at the promoter region of cell type marker genes. **e,** Heat map showing the relative expression of the cell type-specific DEGs. **f**, Heat map showing chromatin accessibility of the cell type-specific DARs.

#### Supplementary Fig.2

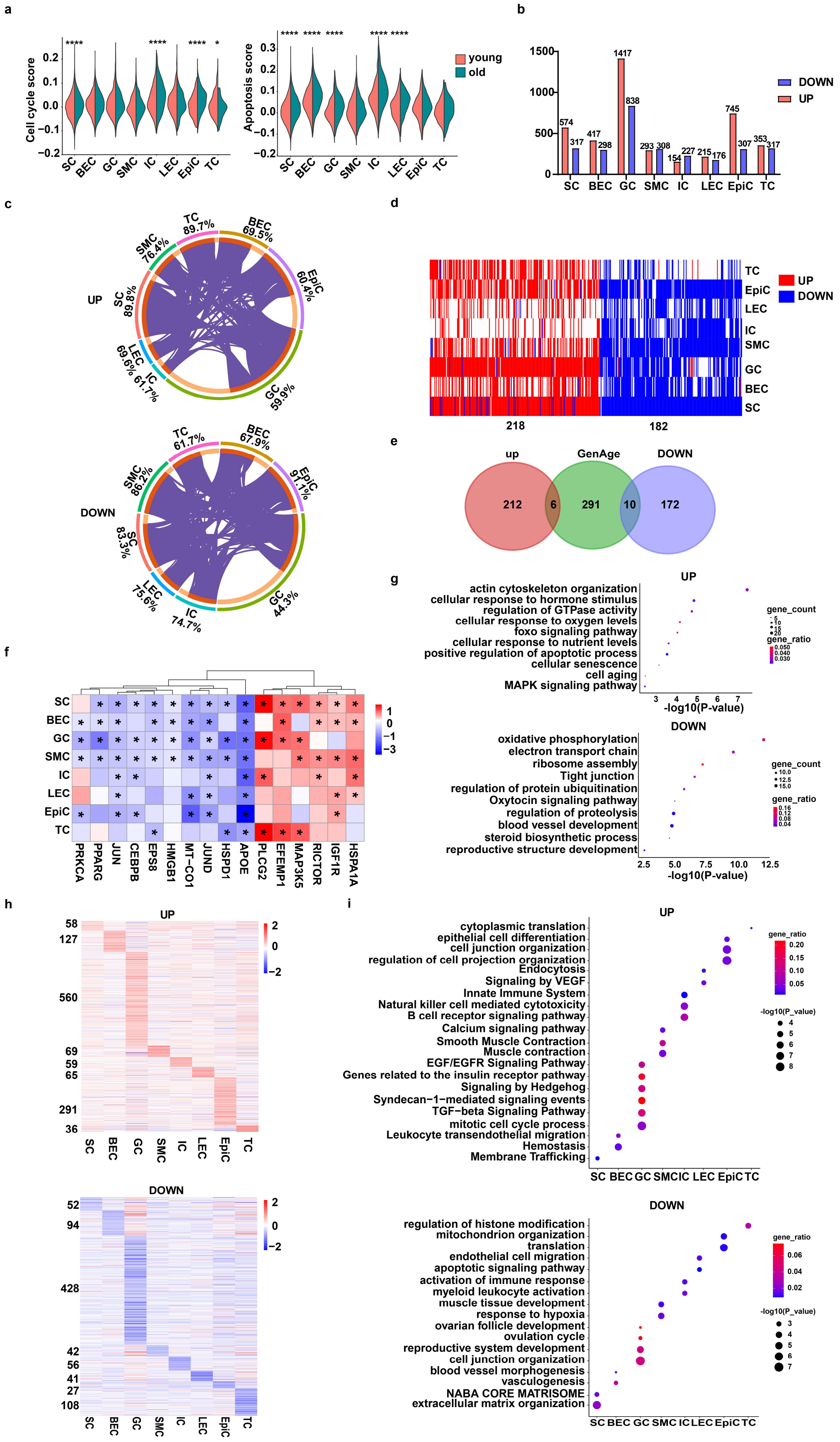

**Supplementary Fig.2: Cell type assignments for human ovary snRNA-seq and snATAC-seq datasets**

**a,** Split violin plots showing the cell cycle score (left) and apoptosis (right) in each cell type from young and aged ovaries. (Two-sided Wilcoxon test; *$P$<0.05, ****$P$<0.0001). **b**, Bar plots depicting the number of ovarian ageing-associated DEGs for each cell type. **c**, Circos plots depicting the overlaps among gene lists of ovarian ageing-associated up-regulated DEGs (left) or down-regulated DEGs (right) for each cell type. The inner-circle represents gene lists, and purple curves link identical genes. The genes that hit multiple lists are colored in dark orange, and genes unique to a list are shown in light orange. **d,** Heat map showing the genes that are differentially expressed in at least 4 cell types between young and aged ovaries. **e,** Venn diagrams of the overlaps between ovarian ageing-associated up- or down-regulated DEGs and human ageing-associated genes from the GenAge database. **f,** Heat map displaying log2 fold changes in gene expression (aged vs. young) of ovarian ageing-associated DGEs in the GenAge database. Asterisk (*) indicates the asterisk indicates a statistically significant difference. ($P_{adj}$ < 0.05). **g**, Representative GO terms of up-regulated common DEGs and down-regulated common DEGs. **h**, Heat map displaying log2 fold changes in gene expression (aged vs. young) of the cell type-specific ovarian ageing-associated up- (top) and down-regulated (bottom) DEGs in each cell type. **i**, Representative GO terms of the cell type-specific ovarian ageing-associated up- (top) and down-regulated (bottom) DEGs in each cell type.

**Supplementary Fig.3**

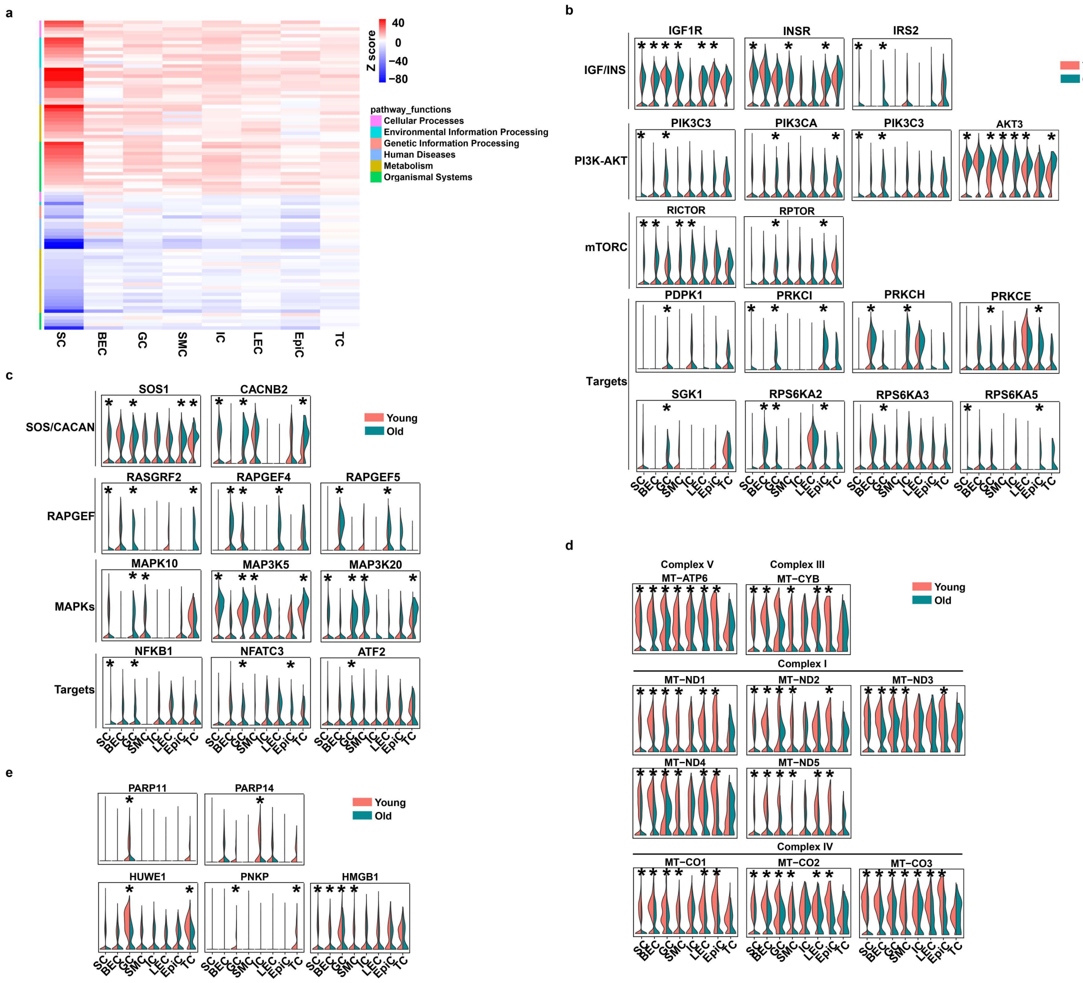


**Supplementary Fig.3: Ageing affects the transcriptional activity of pathways involved in the hallmarks of ageing across cell types**

**a,** Heat map showing all the up- and down-regulated pathways significantly altered in at least 6 cell types during human ovarian ageing ($P_{adj}$<0.05). **b,** Split violin plots showing the expression levels of ovarian ageing-associated DEGs in the mTOR pathway. (MAST test; *$P_{adj}$ <0.05). **c,** Split violin plots showing the expression levels of ovarian ageing-associated DEGs in the MAPK pathway. (MAST test; *$P_{adj}$ <0.05). **d,** Split violin plots showing the expression levels of ovarian ageing-associated DEGs in the oxidative phosphorylation pathway. (MAST test; *$P_{adj}$ <0.05). **e,** Split violin plots showing the expression levels of ovarian ageing-associated DEGs in the base excision repair pathway. (MAST test; *$P_{adj}$ <0.05).

#### Supplementary Fig.4

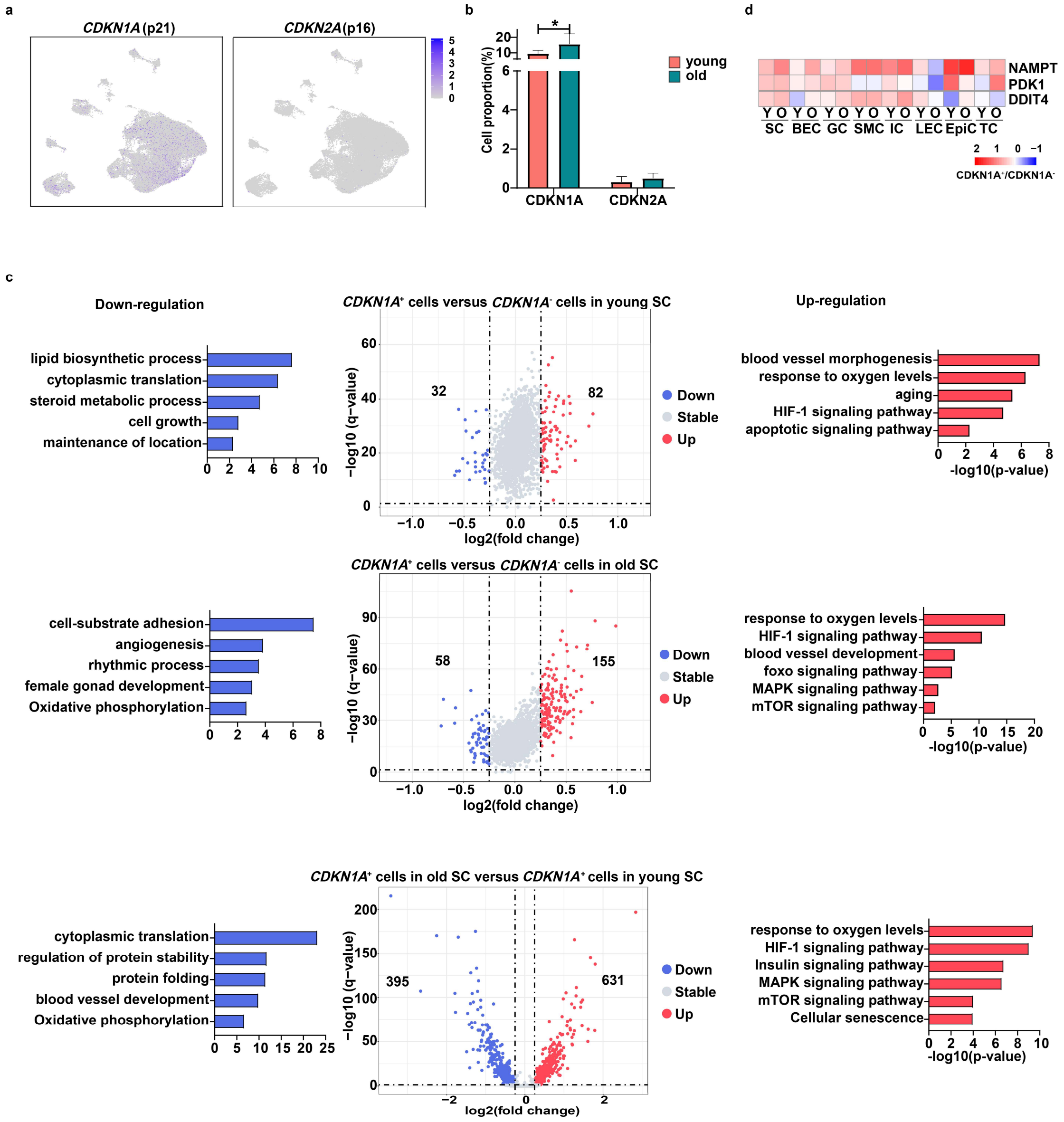


#### Supplementary Fig.4: The proportions and transcriptional programs of *CDKN1A*$^{+}$ cells with age and cell type

**a,** UMAP plots showing the expression of *CDKN1A* (p21) and *CDKN2A* (p16) in the human ovary. **b**, Bar plots represent the proportion of *CDKN1A*$^{+}$ cells and *CDKN2A*$^{+}$ cells in the snRNA-seq data. (Mean±SE; Permutation test; Asterisk (*) indicates FDR<0.05 and abs(log2FD)>1.5). **c,** Middle: Volcano plot depicting differentially expressed genes (DEGs) in *CDKN1A*$^{+}$ cells versus *CDKN1A*$^{-}$ cells from young (top) and old (bottom) stromal cells. Left: Representative GO terms of down-regulated genes in *CDKN1A*$^+$ cells versus *CDKN1A*$^-$ cells from young (top) and aged (bottom) stromal cells, *CDKN1A*$^+$ cells from old stromal cells versus *CDKN1A*$^+$ cells from young stromal cells (middle). Right: Representative GO terms of up-regulated genes in *CDKN1A*$^+$ cells versus *CDKN1A*$^-$ cells from young (top) and aged (bottom) stromal cells, *CDKN1A*$^+$ cells from old stromal cells versus *CDKN1A*$^+$ cells from young stromal cells (middle). **d**, Heat maps displaying log2 fold changes in gene expression (*CDKN1A*$^+$ vs. *CDKN1A*$^-$ cells) of key HIF-1 target genes in each cell type.

Supplementary Fig.5

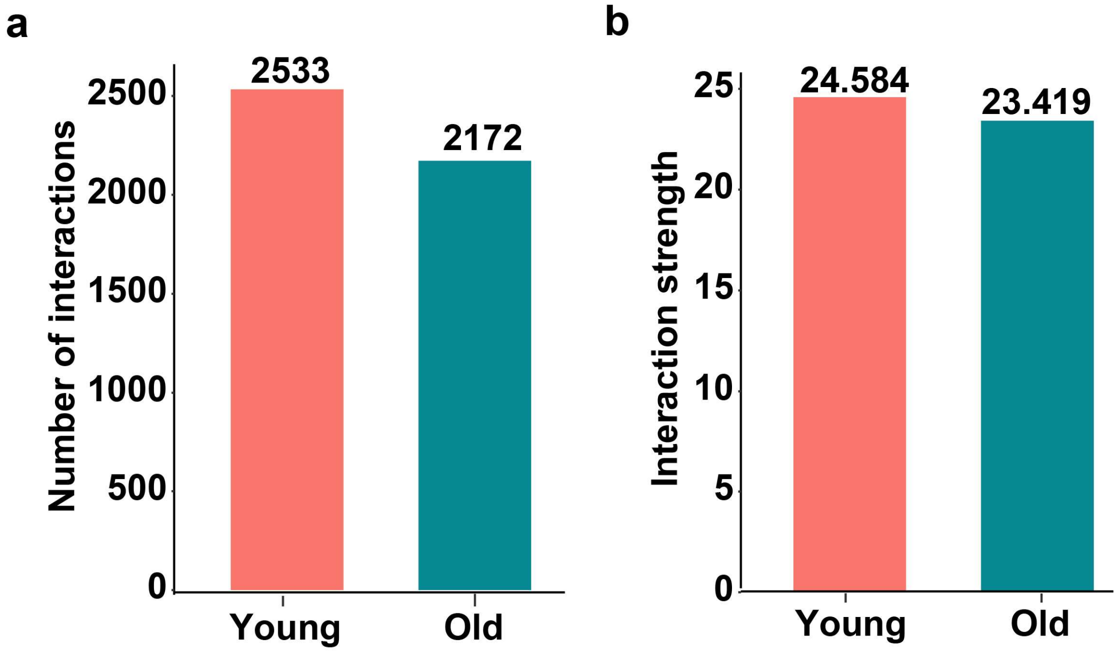


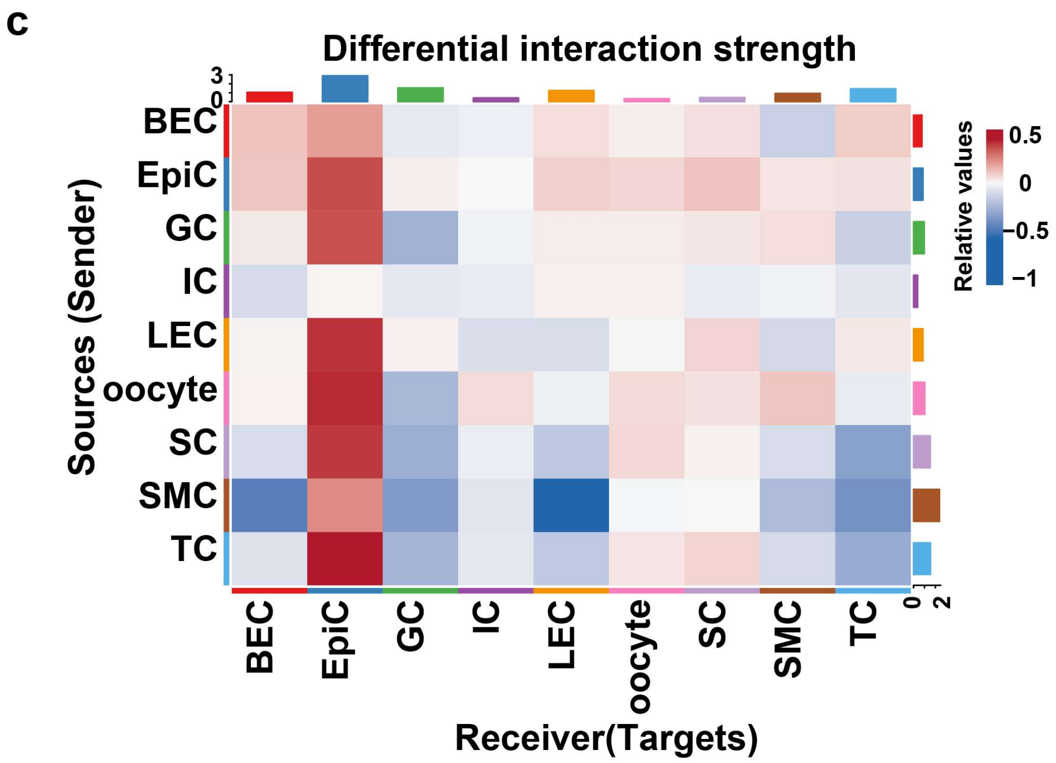


**Supplementary Fig.5: Intercellular interactions between human ovarian cell types during ageing**

**a,** Bar plots represent the number of intercellular interactions in young and old ovaries. **b**, Bar plots represent the strength of intercellular interactions in young and old ovaries. **c,** Heat map of the differential strength of interactions between cell types in young and aged ovaries. The top bar plots represent the sum of each column of values displayed in the heatmap (incoming signaling). The right bar plots represent the sum of each row of values (outgoing signaling).

**Supplementary Fig.6**

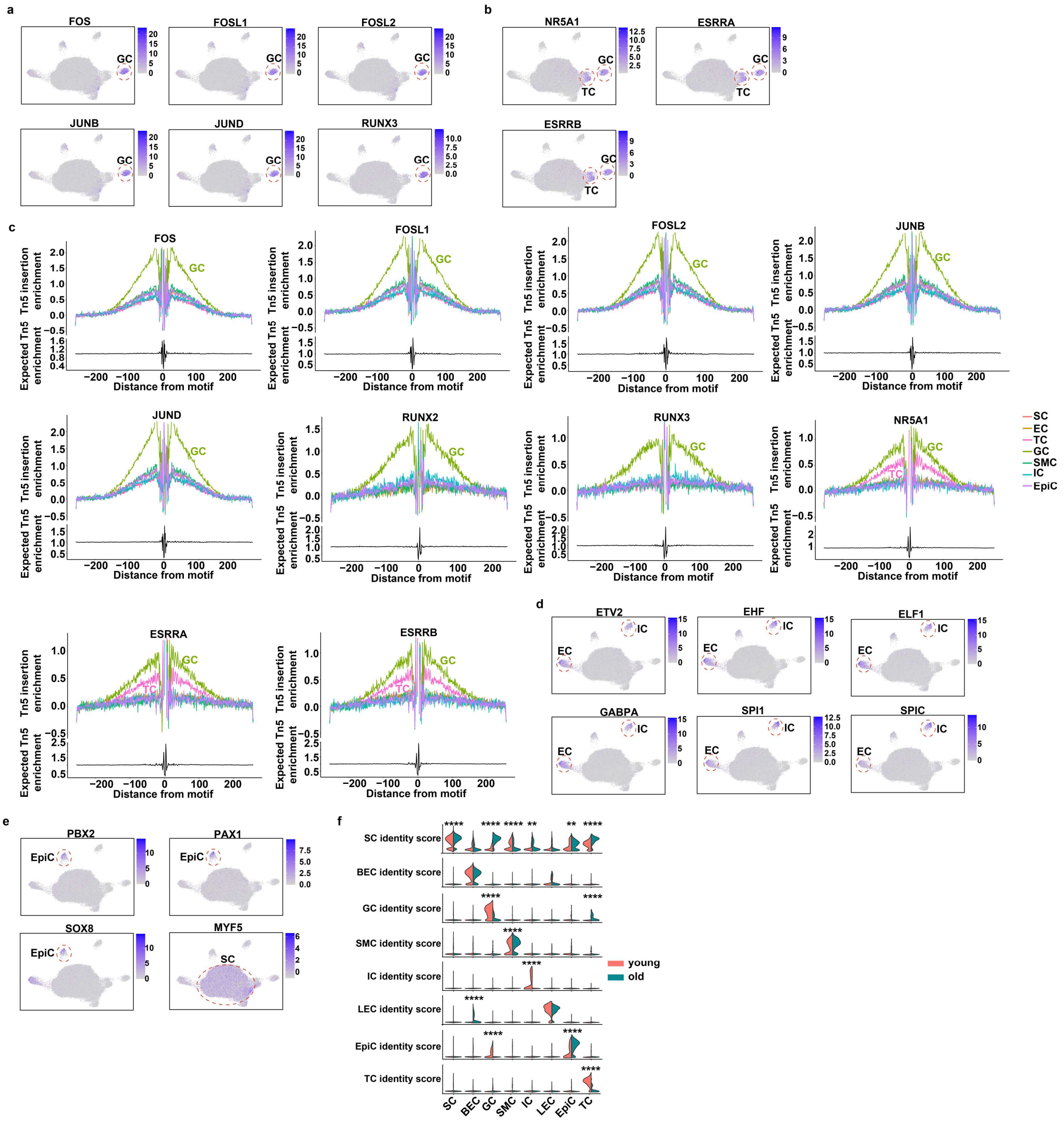


#### Supplementary Fig.6: Cell identity-associated transcription factors in the human ovary

**a**, UMAP plots displaying the chromVAR motif activity of granulosa cell-specific TFs. **b**, UMAP plots displaying the chromVAR motif activity of steroidogenesis-related TFs. **c**, Footprinting analysis of the granulosa cell-specific TFs and steroidogenesis-related TFs across ovarian cell types. **d**, UMAP plots displaying the chromVAR motif activity of endothelial cell and immune cell-specific TFs. **e**, UMAP plots displaying the chromVAR motif activity of epithelial cell and stromal cell-specific TFs. **f**, Split violin plots showing the cell identity score in each cell type from young and aged ovaries. (Two-sided Wilcoxon test; **$P<0.01$, ****$P<0.0001$).

### Supplementary Fig.7

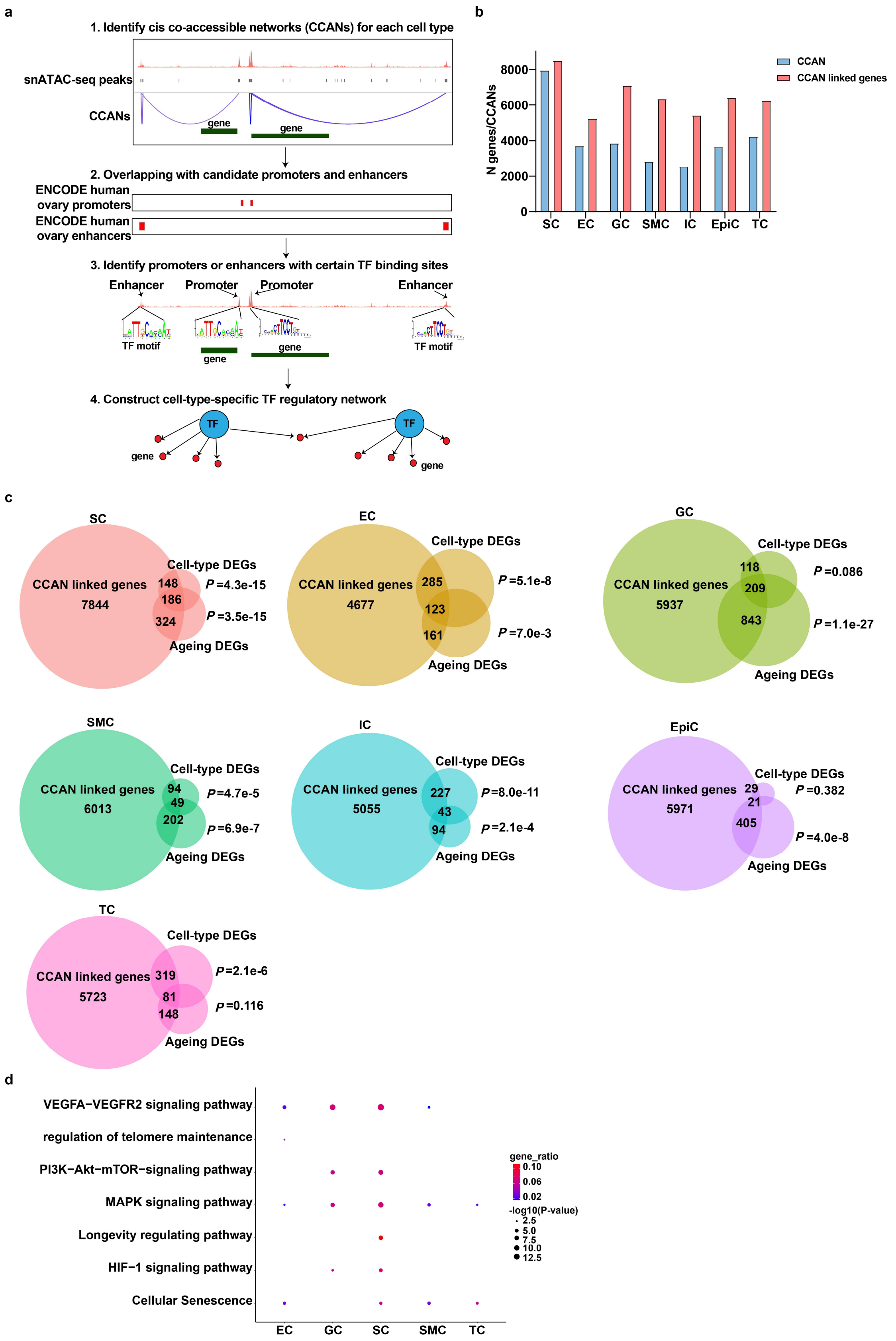

**Supplementary Fig.7: Cell type-specific cis-regulatory networks in the human ovary**

**a**, Schematic of the strategy to construct TF regulatory networks in each cell type. **b**, Bar plots represent the number of CCANs and CCAN-linked genes identified in each cell type. **c**, Venn diagrams of the overlap between CCAN-linked genes and cell type-associated DEGs or ageing-associated DEGs in each cell type. A one-sided Fisher's exact test was used for gene-set overlap significance. **d**, Representative GO terms of CEBPD target genes in each cell type.

#### Supplementary Fig.8

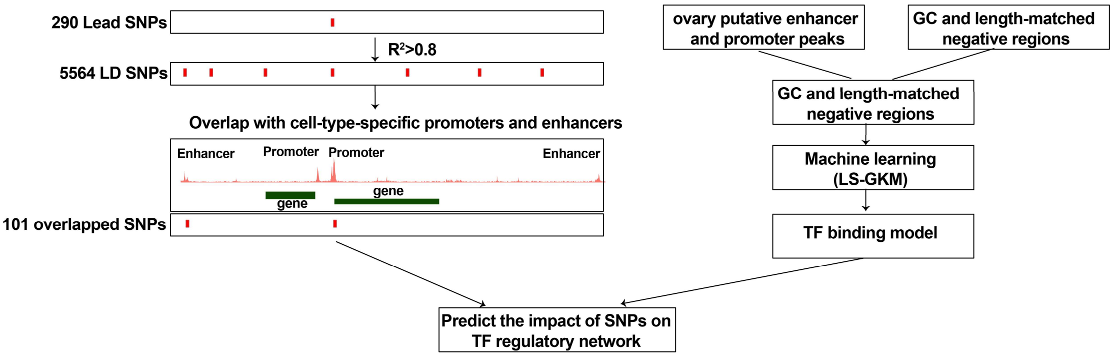


**Supplementary Fig.8: Schematic of the strategy to nominate functional SNPs**

#### Supplementary Fig.9

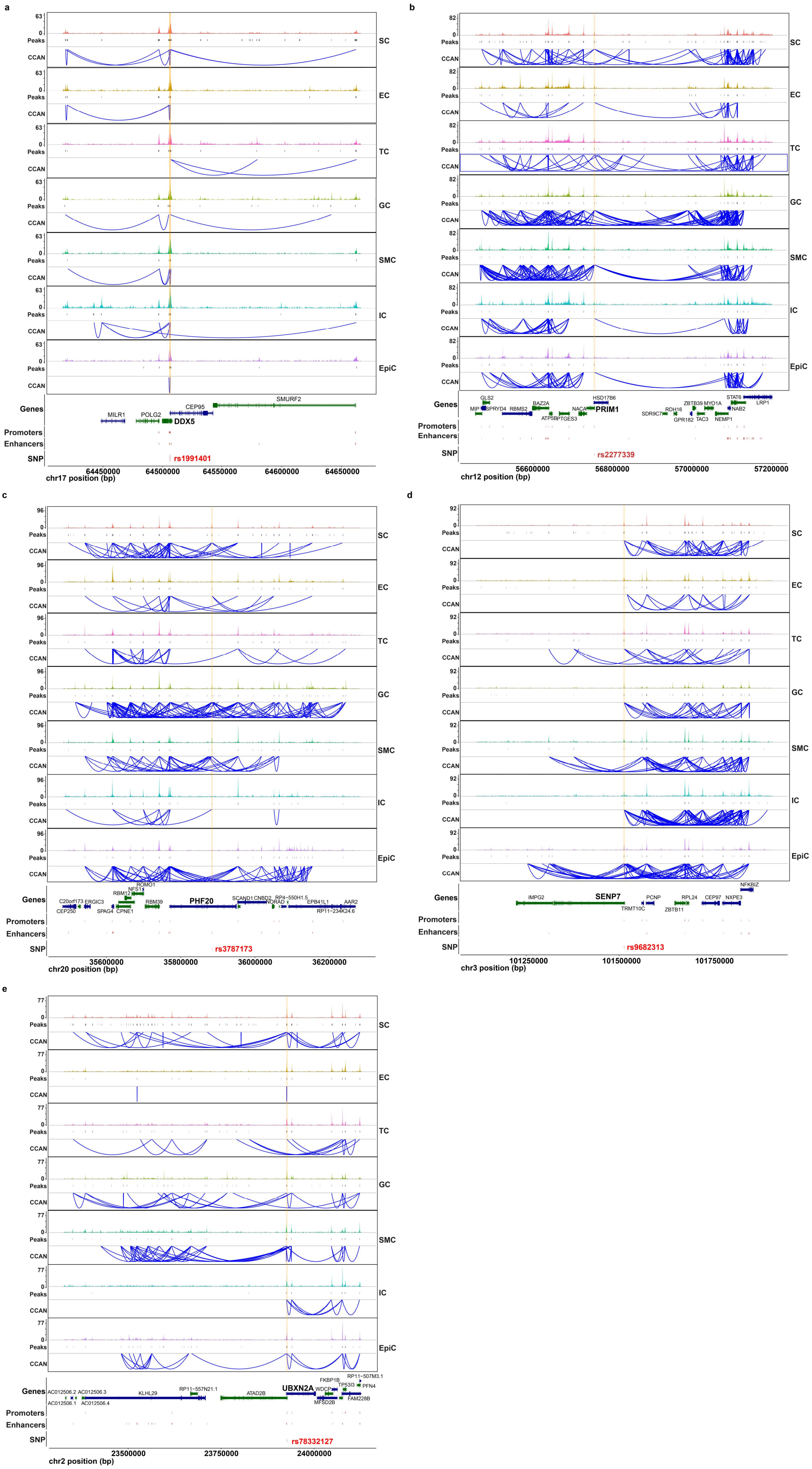

**Supplementary Fig.9: Integration of ANM GWAS and single-nuclei multi-omics identifies the candidate causal variants and cell-type-specific regulatory elements that implicate target genes in the DNA damage response pathway.**

**a-e**, Cis-regulatory architecture at *DDX5* (**a**), *PRIM1* (**b**), *PHF20* (**c**), *SENP7* (**d**), and *UBXN2A* (**e**) gene in each cell type. The snATAC-seq tracks represent the aggregate signals of all cells from a given cell type. The co-accessible peaks inferred by Cicero for each cell type are shown.

**Supplementary Fig.10**

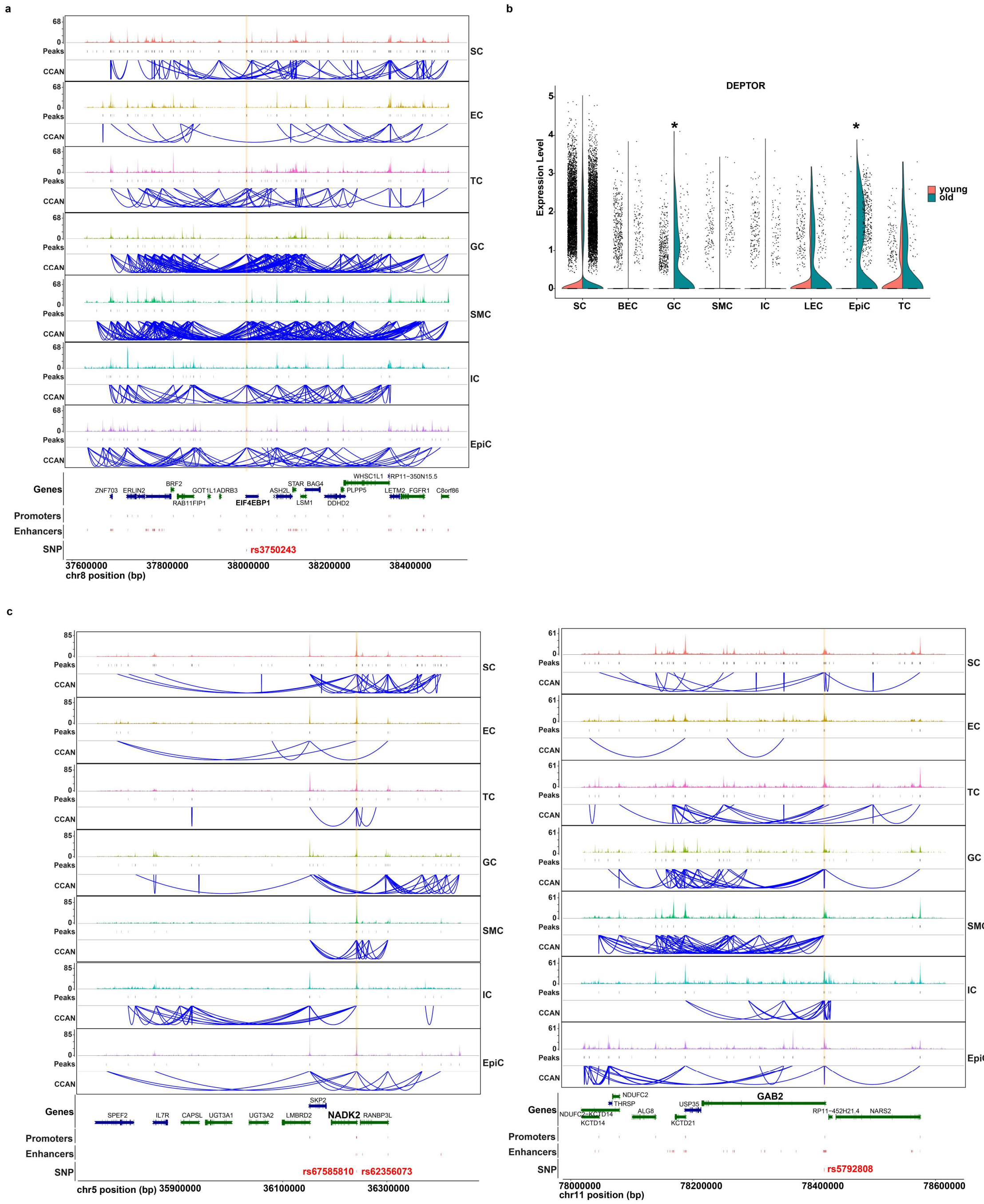


**Supplementary Fig.10: Integration of ANM GWAS and single-nuclei multi-omics identifies the candidate causal variants and cell type-specific regulatory elements that implicate target genes in the mTOR, oxidative phosphorylation, and MAPK signaling pathways.**

**a,c,d**, Cis-regulatory architecture at *EIF4EBP1* (**a**), *NADK2* (**c**), and *GAB2* (**c**) gene in each cell type. The snATAC-seq tracks represent the aggregate signals of all cells from a given cell type. The co-accessible peaks inferred by Cicero for each cell type are shown. b, Split violin plots showing the expression levels of *DEPTOR* in each cell type during ovarian ageing. (MAST test; *$P_{adj}$ <0.05).
